# REV-ERBα mediates complement expression and circadian regulation of microglial synaptic phagocytosis

**DOI:** 10.1101/2020.05.11.088443

**Authors:** Percy Griffin, Patrick W. Sheehan, Julie M. Dimitry, Chun Guo, Michael F. Kanan, Jiyeon Lee, Jinsong Zhang, Erik S. Musiek

**Affiliations:** Department of Neurology, Washington University School of Medicine in St. Louis, St. Louis, MO, USA; Department of Pharmacological and Physiological Science, Saint Louis University School of Medicine, St. Louis, MO, USA; Hope Center for Neurological Disorders, Washington University School of Medicine in St. Louis

## Abstract

The circadian clock has been shown to regulate various aspects of brain health including microglial and astrocyte activation. Here we report that deletion of the master clock protein BMAL1 induces robust increases in the expression of complement genes such as *C3, C4b* and *C1q* in the hippocampus. Loss of downstream REV-ERBα-mediated transcriptional repression led to increases in *C4b* in neurons and astrocytes as well as C3 protein in microglia and astrocytes. REV-ERBα deletion induced complement *C3/C4b* gene expression and increased microglial phagocytosis of synapses in the CA3 region of the hippocampus. Finally, we observed diurnal variation in the degree of microglial synaptic phagocytosis in wild type mice which was abrogated by REV-ERBα deletion. This work uncovers the BMAL1-REV-ERBα axis as a regulator of complement expression and synaptic phagocytosis in the brain, thereby illuminating a novel mechanism of synaptic regulation by the circadian clock.

## INTRODUCTION

The circadian clock orchestrates 24-hour rhythms in various cellular processes through transcriptional-translational feedback loops in most cells of the body (Takahashi, 2017). At the core of the clock’s positive limb is the bHLH-PAS transcription factor BMAL1, which heterodimerizes with CLOCK or NPAS2 to drive the transcription of large sets of clock-controlled genes (Mohawk et al., 2012). BMAL1 transcriptional targets include the negative limb feedback regulator CRY and PER proteins, as well as REV-ERBα and β, nuclear receptors which can also inhibit the actions of the positive limb (Preitner et al., 2002; Retnakaran et al., 1994). Disruption of this circadian machinery is associated with various pathophysiological states including cancer, diabetes and neurodegeneration (Everett and Lazar, 2014; Musiek and Holtzman, 2016; Sulli et al., 2018). Deletion of BMAL1 abrogates circadian clock function and leads to an 85% decrease in REV-ERBα expression in the brain (Musiek et al., 2013). REV-ERBα functions as a transcriptional repressor in many tissues, and has been implicated in regulation of metabolism and inflammation(Everett and Lazar, 2014) Previous work from our group shows that deletion of BMAL1 or its downstream target REV-ERBα causes neuroinflammation and impaired brain functional connectivity (Griffin et al., 2019; Musiek et al., 2013). Diminished BMAL1 and REV-ERBα expression have also been described in mouse models of Alzheimer’s Disease (AD)(Lee et al., 2020; Stevanovic et al., 2017). In AD, memory-associated, synapse-rich regions such as the hippocampus are affected early in the disease course (Braak et al., 2006). Synaptic loss also precedes neuronal death in neurodegeneration (Selkoe, 2002). Circadian dysfunction is also a well-described symptom of AD and other neurodegenerative diseases (Musiek and Holtzman, 2016; Videnovic et al., 2014). Therefore, elucidating the how clock proteins regulate synaptic health is an important step in understanding the connection between circadian dysfunction and neurodegeneration.

A wealth of recent studies have emphasized the critical role of the complement system of the brain in regulating neuroinflammation and synaptic integrity. Synapses labeled with the opsonins C1q and C3 (Stevens et al., 2007) were first described to be pruned by microglia during development (Schafer et al., 2012). C4 protein, encoded by the mouse *C4b* gene, also contributes to synaptic pruning by microglia in vivo (Sekar et al., 2016). Complement-dependent microglial synaptic pruning has also been implicated in the pathogenesis of neurodegenerative and neuropsychiatric diseases (Hong et al., 2016; Litvinchuk et al., 2018; Sekar et al., 2016; Werneburg et al., 2020). Microglial activation is subject to circadian regulation (Fonken et al., 2015; Hayashi et al., 2013), and we have previously described that deletion of BMAL1 or REV-ERBα can induce microglial activation (Griffin et al., 2019; Musiek et al., 2013). Given the roles of the clock in neurodegeneration and microglial activation, we explored a potential role of the core clock in regulating synaptic health. Herein we establish a novel link between the BMAL1-REV-ERBα axis, complement expression, and microglial synaptic pruning in the hippocampus.

## RESULTS

### Disruption of the BMAL1-REV-ERBα axis induces complement upregulation in multiple brain cell types

While analyzing our previously published transcriptomic dataset from Bmal1 KO (BMKO) hippocampal tissue, we observed a striking upregulation of several complement transcripts, in particular *C4b* and *C3*, two genes which are critical for synaptic phagocytosis (Fig 1A). Other complement related transcripts including *C1qc, C1qb, C1qa C1ra*, and *C1rb* were also significantly increased in BMKO hippocampus (Fig 1A). *C4b* was similarly increased in cerebral cortex samples from 4mo tamoxifen-inducible global BMAL1 KO mice (CAG-Cre^ERT2^;Bmal1^f/f^) in which *Bmal1* was deleted at 2mo, demonstrating that this is not a developmental phenomenon (Fig S1).

**Figure 1:**
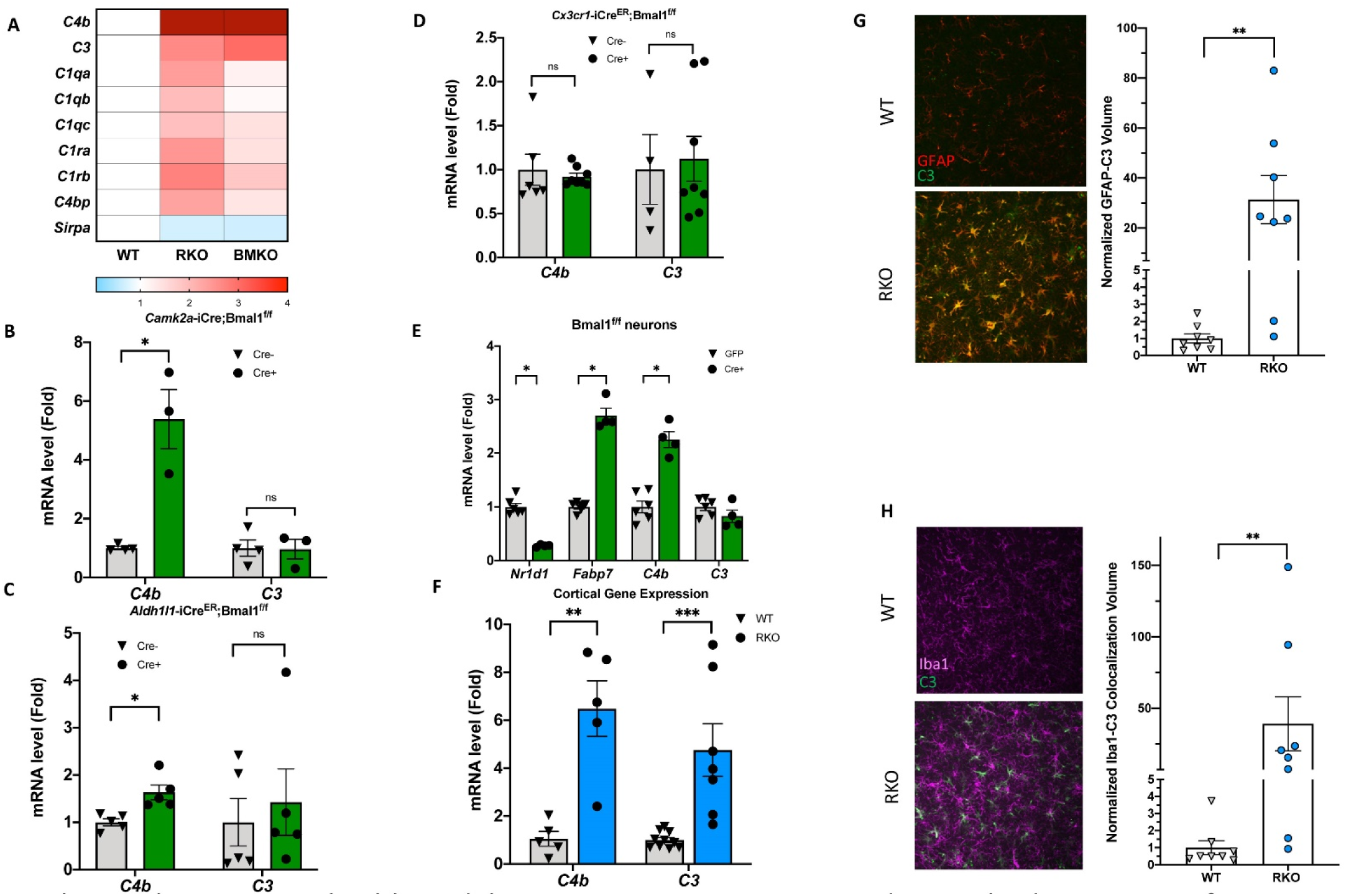
Rev-erbα regulates complement expression in multiple cell types downstream of BMAL1. A. Relative expression of complement related transcripts taken from microarray analysis performed on hippocampus from 5mo RKO and littermate WT mice (N= 3/genotype) as well as BMAL1 KO mice (N=2). B. qPCR analysis of 11month old control (Cre-) and neuron-specific *Bmal1 KO* mice (*Camk2a-iCre*+;*Bmal1*^*fl/fl*^) for complement genes (N=3-4/group). C. qPCR analysis of control (Cre-) and astrocyte-specific *Bmal1 KO* mice (Aldh1l1-Cre^ERT2^+;Bmal1^fl/fl^) for complement genes (N = 5 mice/group). D. qPCR analysis of control (Cre-) and microglia-specific Bmal1 KO mice (Cx3cr1-Cre^ERT2^+;Bmal1^fl/fl^) for complement genes (N = 4-8/group). For C and D, all mice (Cre- or +) were treated with tamoxifen at 2mo and harvested at 4mo, mixed sexes, Cre-littermates were used as controls. E. qPCR analysis of mRNA from primary cortical neurons isolated from Bmal1^fl/fl^ mice, treated with AAV8-GFP (control) or AAV8-Cre (n = 4-5/group). F. qPCR analysis of WT or RKO mouse cortical tissue for complement genes (N= 5-10 mice/group). G. Representative 40X maximum intensity projections of hippocampus of 5mo WT or RKO mice stained for C3 and GFAP as well as the associated normalized volumes for C3-GFAP staining (n=8 mice, N=4/group). H. Representative 40X maximum intensity projections of 5mo WT or RKO mice stained for C3 and Iba1 as well as the associated normalized volumes for C3-Iba1 staining (n=8, N=4/group). In G and H, each point represents the average of 3 sections from a single mouse. *p<0.05 **p<0.01 ***p < 0.001 by 2-tailed T-test with Welch’s correction.

To determine the cell type(s) in which BMAL1 deletion induces complement gene expression, we examined cerebral cortex tissue from pan-neuron-(CamK2a-iCre;Bmal1^fl/fl^)((Izumo et al., 2014), astrocyte-(Aldh1l1-Cre^ERT2^;Bmal1^fl/fl^)(Lananna et al., 2018) and microglia-specific (Cx3cr1-Cre^ERT2^;Bmal1^fl/fl^)(Parkhurst et al., 2013) BMAL1 knockout mice. Both astrocyte- and microglia-specific Bmal1 KO mice were treated with tamoxifen at 2mo and harvested 2 months later. Notably, *C4b* mRNA was strongly induced in the neuron-specific BMAL1 KO mice, while *C3* was not (Fig 1B). *C4b* was also induced in astrocyte-specific BMAL1 KO mice (Fig 1C). *C4b*, but not *C3*, was induced in global inducible Bmal1 KO mice (CAG-Cre^ERT2^;Bmal1^f/f^) 2mo after gene deletion (Fig. S1) Under basal conditions, neither *C4b* nor *C3* was induced in microglia specific BMAL1 KO mice (Fig 1D). Deletion of BMAL1 in primary Bmal1^fl/fl^ neuron cultures via infection with an AAV8-Cre viral vector (versus AAV8-eGFP control) suppressed the BMAL1 transcriptional target REV-ERBα (*Nr1d1*) by 85% and induced *C4b* expression but caused no increase in *C3* (Fig 1E). BMAL1 deletion also increased expression of *Fabp7* (Fig 1E), a known target of REV-ERBα-mediated transcriptional repression (Schnell et al., 2014). This suggested that the upregulation of *C4b* gene expression observed with loss of BMAL1 could be mediated by de-repression as consequence of downstream REV-ERBα loss. Accordingly, global deletion of REV-ERBα caused striking increases in *C4b, C3* and other complement transcripts in the hippocampus as assessed by microarray analysis (Fig 1A) and confirmed in separate samples by qPCR (Fig 1F). Increased C3 protein expression was observed in both activated astrocytes (Fig 1G) and microglia (Fig 1H) in the hippocampus of 5mo REV-ERBα KO (RKO) mice. The finding that *Bmal1* deletion induces *C4b* mRNA early, but not *C3*, but that C3 mRNA and protein are increased at later ages, suggest that REV-ERBα directly represses *C4b* expression in neurons and astrocytes, but the induction of *C3* in both BMKO and RKO brain is likely secondary to inflammatory glial activation which occurs over time.

### REV-ERBα regulates microglial synaptic engulfment

We previously demonstrated that global REV-ERBα deletion induced microglial activation in vivo (Griffin et al., 2019). Given those results and the observation of increased *C4b* and *C3* expression in RKO brains, we examined the possibility that these changes would enhance synaptic phagocytosis in 4-6mo RKO mice. We primarily focused on the mossy fiber synapses in the CA2/3 region of the hippocampus, as these large synapses (thorny excrescences) can easily be stained and imaged using standard confocal microscopy. Triple-labeling of tissue sections was performed with antibodies against synaptophysin (a marker of presynaptic neuronal terminals – Fig 2A), CD68 (a microglial lysosome marker – Fig 2B) and Iba1 (to define microglial cell bodies and processes – Fig 2C). CD68 was used to ensure that the colocalized synaptic material was actually within the microglia phagosome. 3D reconstructions were made and total volumes of engulfed synaptic material were calculated. We noted a 10-fold increase in engulfed synaptic material in the hippocampus of RKO mice compared to their WT littermates at 6mo (Fig 2F). In the RKO microglia, we observed synaptic material in the microglial process and cell body (Fig 2Eii, Eiii), whereas WT microglia only had engulfed synaptic material in the cell body (Fig 2Ei).

**Figure 2:**
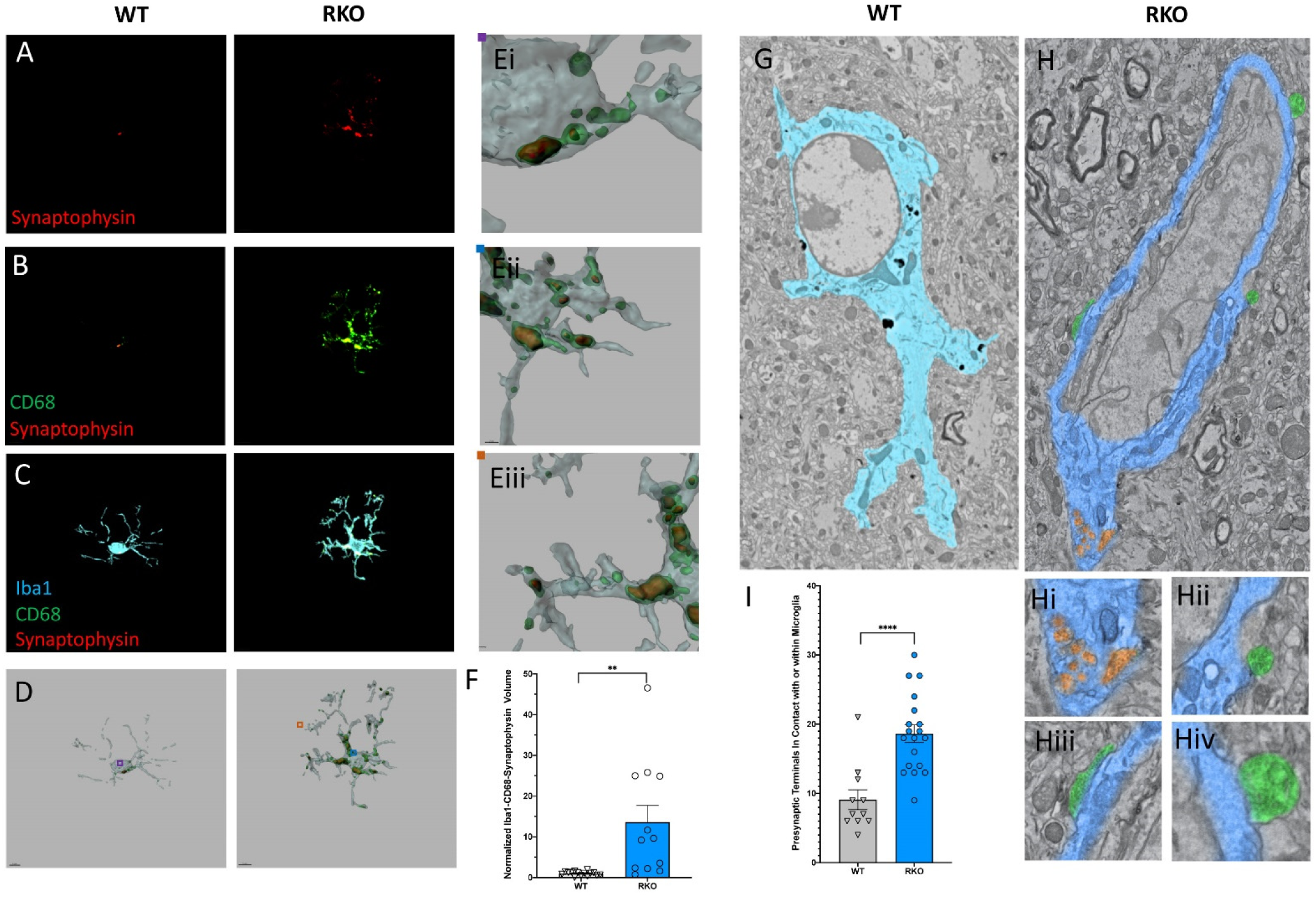
Rev-erbα deletion increases synaptic phagocytosis in the CA3 region of the hippocampus. A. Image of synaptophysin from microglia in the CA3 of 4-6mo WT or RKO mice sacrificed at 11AM. B. Colocalized synaptophysin/CD68 staining from microglia in the CA3 of 4-6mo WT or RKO mice sacrificed at 11AM. C. Representative images of Individual microglia from CA3 stained with Iba1, CD68, and synaptophysin from 4-6mo WT or RKO mice sacrificed at 11AM. D. Representative 3D surface rendering of microglia showing engulfed presynaptic material in lysosomes with zoomed in lysosomal content from WT mice in Ei and littermate RKO mice in Eii/Eiii. F. Quantification of the normalized Iba1-CD68-Synaptophysin volumes from microglia in the CA3 region of the hippocampus of 4-6mo WT and RKO mice (each point is average of 3 sections from one mouse, N = 15 and 12 RKO mice). G. Annotated, representative scanning electron micrographs of microglia in the CA3 of WT (Cyan) or RKO (Royal blue) mice sacrificed at 11AM with zoomed in pictures of engulfed presynaptic terminals in Hi and presynaptic terminals in contact with microglia in Hii, Hiii and Hiv. I. Quantification of presynaptic terminals in contact with or engulfed by microglia in the CA3 of WT or RKO mice. Each point represents one field of view, N=2 mice/genotype. **p < 0.01,**** p<0.0001 by 2-tailed T-test with Welch’s correction.

To corroborate these results, we also performed large area scanning electron microscopy (SEM) experiments of the CA3 region of 6mo WT and RKO mice. Using this method, we visually confirmed an increased number of presynaptic terminals within or in contact with microglia in the hippocampus of RKO mice compared to WT (Fig 2G, 2H, 2H(i-v), 2I). Interestingly, in our RKO mice we noted a downregulation in the expression of the *Sirpa* gene, which codes for the protein SIRPα (Fig 1A). SIRPα was recently described as a surface receptor on microglia that serves as a “do-not-eat-me” signal (Lehrman et al., 2018). Taken together, our results suggest that REV-ERBα deletion induces changes in microglial signaling that precipitates synaptic engulfment.

### CA3 synapses are reduced by REV-ERBα deletion

Following our observations of synaptic pruning in the RKO mice, we investigated the status of the synapses in the CA3 region of the hippocampus. Synapses were double labeled by using the presynaptic marker synaptophysin and the postsynaptic marker homer1. In RKO mice, we observed a significant decrease in the synaptic volume in the CA3 region of the hippocampus by synaptophysin staining (Fig 3A), homer1 staining (Fig 3B) and their colocalization (Fig 3C) compared to their WT littermates. To further confirm these results, we counted synapses in large area SEM images from the stratum lucidum of WT or RKO mice. Again, we noted a decrease in the number of synapses in the RKO mouse CA3 by SEM, as compared to their WT littermates (Fig 3D).

**Figure 3:**
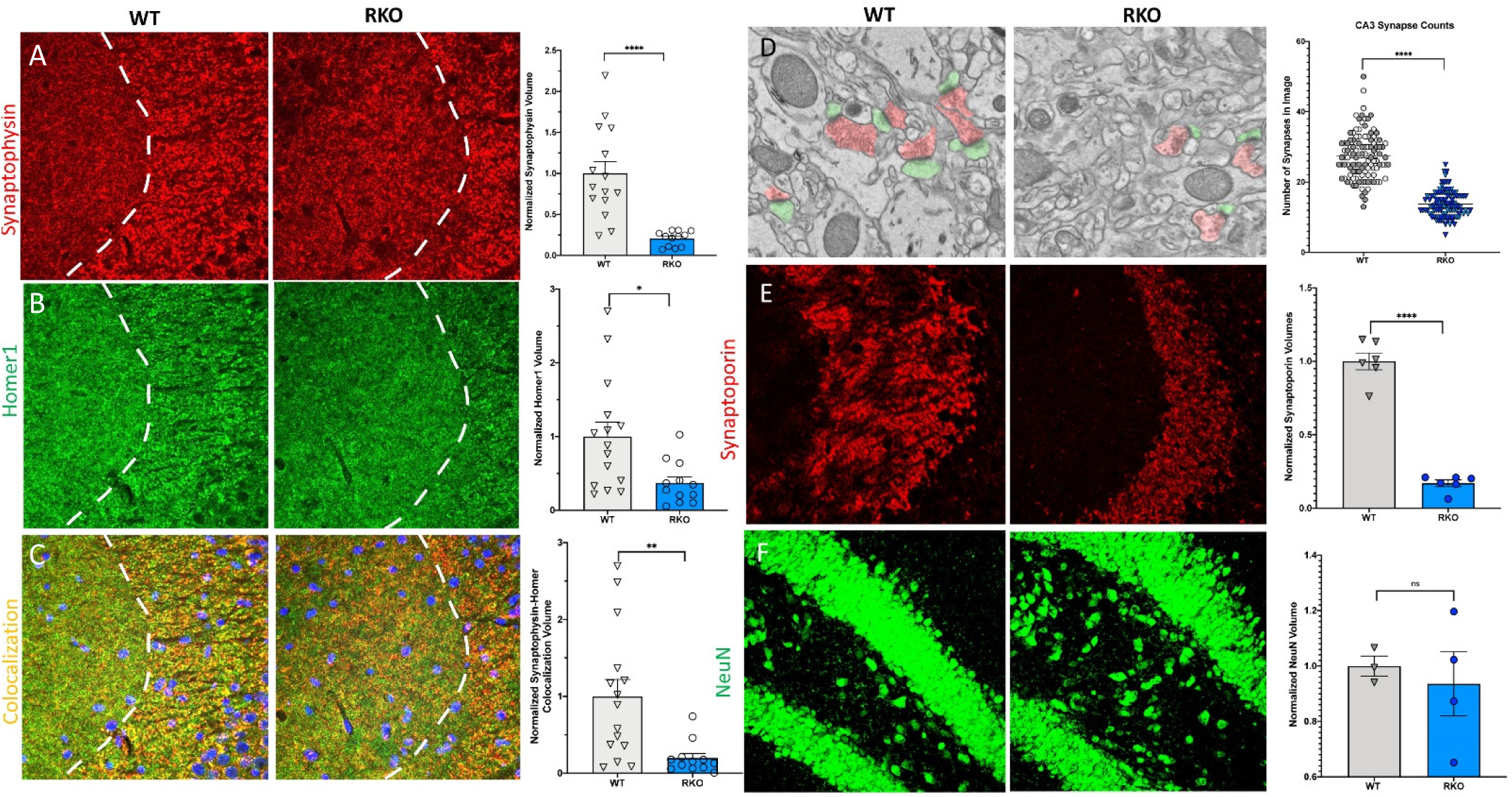
Rev-erbα deletion induces synapse loss in the CA3 region without neuronal loss in the dentate gyrus. A. 60X representative maximum intensity projections showing synaptophysin staining in the CA3 of 4-6mo WT and RKO mice with the associated normalized volume quantification (N = 15 WT and 12 RKO mice). B. 60X representative maximum intensity projections showing homer1 staining in CA3 of 4-6mo WT and RKO mice with the associated normalized volume quantification. C. 60X representative maximum intensity projections showing colocalized synaptophysin and homer1 staining in CA3 of 4-6mo WT and RKO mice with the associated normalized volume quantification. D. Representative scanning electron micrographs of synapses (presynaptic terminal in red, postsynaptic terminal in green) as well as the associated synapse counts for 4-6mo WT and RKO mice. Each point represent one field of view, N=2 mice/genotype, shading indicates each mouse). E. 60X representative maximum intensity projections showing synaptoporin staining in the CA3 of 4-6mo WT and RKO mice with the associated normalized volume quantification (N=6/genotype). F. 40X representative maximum intensity projections showing NeuN staining in the dentate gyrus of 4-6mo WT and RKO mice with the associated normalized volume quantification (N = 3-4 mice/genotype). *p<0.05, **p < 0.01, **** p<0.0001 by 2-tailed T-test with Welch’s correction. In all panels except D, each point represent the average of three sections from a single mouse.

To ensure that our observations were not due purely to changes in synaptophysin protein expression in the presynaptic terminal, synapses were also stained with a second presynaptic marker, synaptoporin, and quantified. Synaptoporin was used because it is enriched in the mossy fiber synapses of the hippocampus (Singec et al., 2002). Again, we observed a similar decrease in synaptic volume in CA3 of RKO mice compared to their WT littermates using synaptoporin staining (Fig 3E). Interestingly, we did not observe significant differences in synaptic volumes between WT and RKO mice in the CA1 region of the hippocampus (Fig S2A-C), suggesting that some terminals may be more susceptible to loss than others. To determine whether the loss in synapses was due to a loss of neuronal cell bodies, we quantified the volume of neuronal nuclei of the dentate gyrus, which project to CA3, via NeuN staining. We found no significant difference between the neuronal nuclear volumes of the dentate gyrus between the WT and RKO mice (Fig 3F). Taken together, our data suggest that deletion of REV-ERBα results in robust synaptic loss in the CA3 region without obvious neuronal loss.

### Time-of-day variation in microglial phagocytosis is regulated by REV-ERBα

In the brain, REV-ERBα displays daily oscillation on the mRNA level – with its peak at zeitgeber time (ZT) 8 (2pm) and its trough at ZT20 (2am) (Chung et al., 2014). We have also previously described oscillations in microglial activation, with low activation at ZT5 and high activation at ZT17 (Griffin et al., 2019). Consequently, we investigated microglial synaptic engulfment at different times of day. WT(C57BL/6) mice sacrificed at ZT5 (11AM) or ZT17 (11PM), and hippocampal sections were triple labeled with Iba1, CD68, and synaptophysin. Microglia from the CA3 region of the hippocampus in mice sacrificed at ZT17 showed significantly more engulfed presynaptic protein than those at ZT5 (Fig 4A, Fig 4B, Fig 4Bi, Fig 4C). To further establish these findings, we used large area SEM to count the number of presynaptic terminals in contact with or within microglia at ZT5 or ZT22. In that experiment, we also noted a higher number of presynaptic terminals in contact with and within the microglia at ZT17 (Fig 4D, Fig 4E, Fig 4Ei, Fig 4F). As a final confirmation of this finding, we injected CX3CR1^GFP^ mice, which express GFP in microglia (Jung et al., 2000), with an AAV-CaMKII-mCherry viral vector in the retrospenial cortex. 4 weeks later, we collected brain samples at ZT5 and 17, and calculated the volume of mCherry+ material in individual GFP+ microglia. Here, we found that despite interactions between microglia and neurons at both timepoints, microglia at 11PM (CT17) engulfed more mCherry than those at 11AM (ZT5) (Fig 4G-I). RKO mice harvested at ZT5 and 17 showed a persistently increased level of synaptic engulfment in microglia, but with no time-of-day variation (Fig 4J), suggesting that daily oscillations in REV-ERBα may mediate rhythms in microglial synaptic phagocytosis. Overall, our data establish a REV-ERBα-dependent rhythm for the engulfment of neuronal materials in the brain parenchyma by microglia.

**Figure 4:**
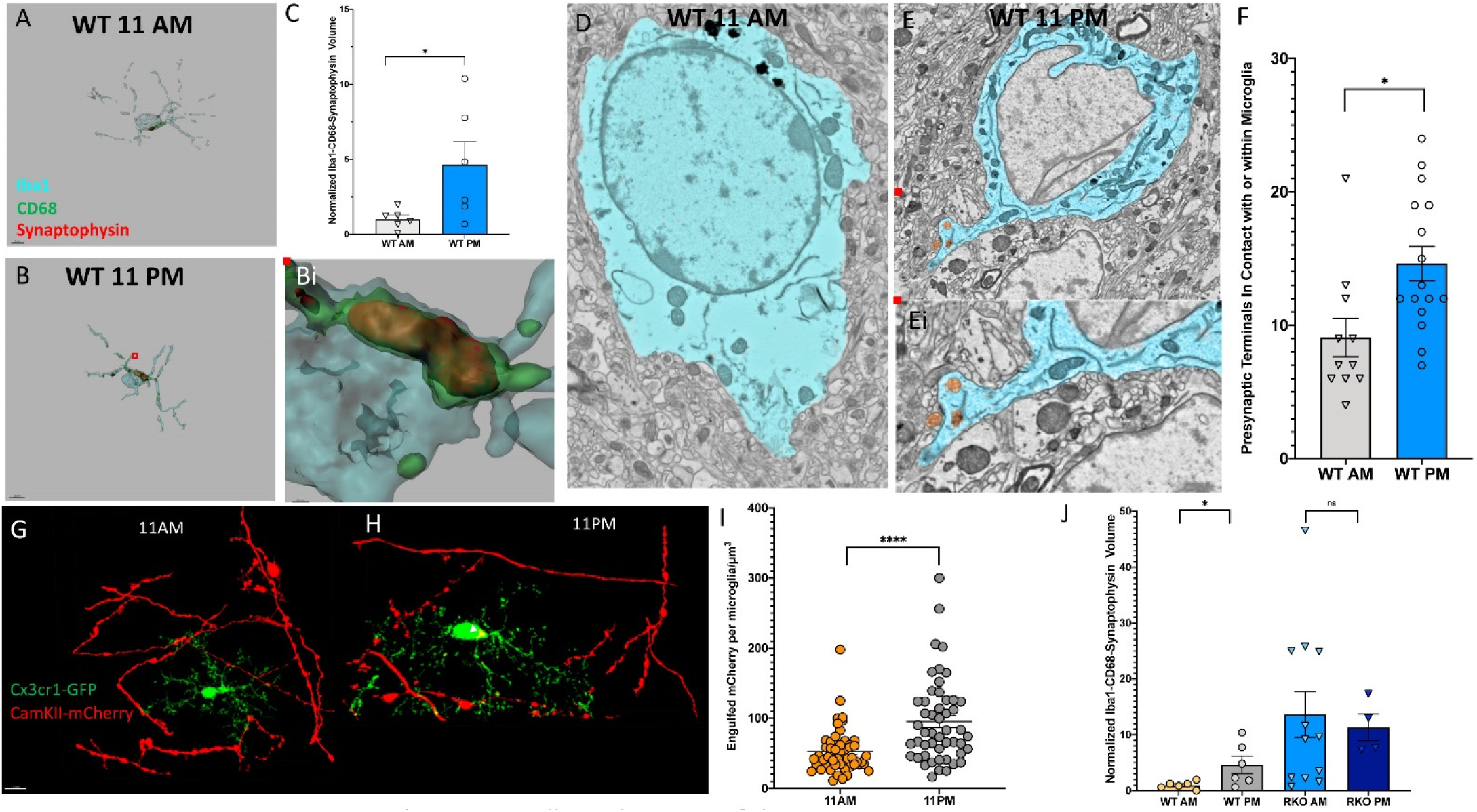
Microglial phagocytosis oscillates. 3D surface rendering of Iba1, CD68 and synaptophysin in microglia from 6 mo WT mice sacrificed at A. 11AM or B. 11PM. Bi. Enlarged rendering showing engulfed synaptophysin in microglia from WT mice sacrificed at 11PM. C. The normalized volume of Iba1-CD68-Synaptophysin showing microglial phagocytosis of synapses (N=5 mice/timepoint). Representative scanning electron micrographs of microglia in the CA3 of WT mice sacrificed at D. 11AM or E. 11PM as well as Ei.a zoomed in picture of presynaptic terminals within the microglia. F. Quantification of presynaptic terminals in contact with or engulfed by microglia in WT mice sacrificed at 11AM or 11PM. Each point represent one field of view, N=2 mice/timepoint). Representative 100X images of microglia and neuronal axons from 6-8mo CX3CR1^GFP^ mice injected with an AAV2-CamKII-mCherry and sacrificed at G. 11AM or H. 11PM as well as the I. associated quantification of the engulfed mCherry volume per microglia. (n=50 microglia counted/mouse, N=2 mice/genotype) J. Normalized volume of Iba1-CD68-Synaptophysin for WT or RKO mice sacrificed at 11AM or 11PM. (N = 4-12/group) *p < 0.05, ****p<0.0001 by 2-tailed T-test with Welch’s correction. In panels C and J, each point represent the average of three sections from a single mouse.

## DISCUSSION

The current study shows that loss of the circadian protein BMAL1 causes upregulation of complement gene *C4b* in neurons and astrocytes, as well as increase astrocytic *C3* expression, all via loss of downstream REV-ERBα-mediated transcriptional repression. Deletion of REV-ERBα leads to microglial activation, increase *C4b* and *C3* expression, and increased microglial synaptic phagocytosis in CA3 region of the hippocampus. Finally, we demonstrate a time-of-day variation in synaptic phagocytosis in the hippocampus of WT mice, which is lost after REV-ERBα deletion. Our finding suggests that the BMAL1-REV-ERBα axis regulates daily rhythms in synaptic phagocytosis, and that loss of REV-ERBα de-represses complement gene expression and locks the brain in a pro-phagocytic state.

The mechanisms by which *Bmal1* deletion leads to increased complement gene expression are complex and multicellular but appear to depend on REV-ERBα. REV-ERBα expression is decreased by ∼85% following *Bmal1* deletion, and REV-ERBα deletion phenocopies the complement gene expression increases seen in *Bmal1* KO brain. Indeed, it is well established that REV-ERBα functions as a transcriptional repressor (Harding and Lazar, 1995). Loss of REV-ERBα-mediated repression is a general mechanisms governing increased transcript expression following *Bmal1* deletion - as evidence by strong upregulation of the REV-ERBα repression target *Fabp7* following *Bmal1* deletion (Schnell et al., 2014). Notably, we queried an existing REV-ERBα ChIP-seq database from brain tissue (GEO Series GSE67962, Zhang et al., 2015), and did not find any peaks corresponding to *C4b* or *C3*. However, REV-ERBα is known to regulate transcription not only by direct binding to RORE sites, but also by regulating enhancer function and eRNA expression to alter gene expression in *trans* (Lam et al., 2013, and by altering chromatin looping(Kim et al., 2018). Thus, it is likely that REV-ERBα regulates *C4b* expression in one of these alternative ways which is not apparent by typical ChIP-seq. Tissue-specific deletion of Bmal1 shows that this Bmal1-REV-ERBα axis controls *C4b* expression in neurons and astrocytes, but less so in microglia. Recent single-nucleus RNAseq data suggests that *C4b* is prominently expressed in oligodendrocytes (Zhou et al., 2020). However, we have not yet evaluated the effect of BMAL1 or REV-ERBα deletion on *C4b* in this cell type. Presumably, *C4b* expression is low in neurons and astrocytes under normal conditions but is increased by *Bmal1* or REV-ERBα deletion. Work from the Stevens and Carroll labs identified *C4b* expression in neurons which then tagged synapses and facilitated synaptic pruning, and showed that deletion of *C4b* prevent accumulation of C3 protein at synapses, as C4 protein in upstream of C3 activation in the classical complement pathway (Sekar et al., 2016).

*Bmal1* KO and RKO mice at 5-6mo also show an increase in *C3* transcript and C3 protein in astrocytes. However, Tissue-specific *Bmal1* deletion in neurons or astrocytes causes pronounced *C4b* increases but no *C3* increases, suggesting that C3 expression in astrocytes and microglia is a secondary response to increased neuroinflammation in aged global Bmal1 KO or RKO mouse brain. Global inducible *Bmal1* KO mice also do not show *C3* induction at 2mo post-tamoxifen (despite high *C4b* expression), as these mice have not developed a full neuroinflammatory response at that age. Our data suggests that full induction of C3 in *Bmal1* and RKO brain requires a time-dependent inflammatory response. We previously showed that REV-ERBα deletion increased *Traf2* expression and increased NFκB signaling (Griffin et al., 2019). NFκB signaling has also been shown to increase C3 expression and release from astrocytes (Lian et al., 2015). Thus, REV-ERBα loss could induce C3 in astrocytes through the induction of NFκB signaling. We also noted an increase in *C1q* expression in RKO primary neurons, microglia and whole hippocampus. In the classical complement cascade, C1 and C4 are upstream of C3 (Zipfel and Skerka, 2009). Functionally, C4 loss is associated with less synaptic tagging with C3 and thus less synaptic pruning (Sekar et al., 2016), suggesting that C4 expression can drive C3-dependent synapse pruning. Additionally, we noted that REV-ERBα regulated the “do-not-eat-me” signal *Sirpa* (encoding SIRPα) in our microarray data. Loss of inhibitory signaling from SIRPα makes microglia more likely to prune synapses, since SIRPα is primarily expressed on microglia in the CNS (Zhang et al., 2014). Therefore, our data suggest that de-repression of *C4b* in neurons and *Traf2* in astrocytes and microglia, as well as diminished *Sirpa* expression in microglia, lead to a “perfect storm” of complement expression and microglial activation that promotes synaptic phagocytosis.

We focused on the CA2/3 mossy fiber synaptic boutons because they are large and easily labeled with 2 synaptic vesicle markers – synaptoporin and synaptophysin (Grosse et al., 1998). However, we did not observe synapse loss in the CA1 region, which may represent a regional variability in BMAL1-REV-ERBα mediated synaptic pruning. Certainly, it is possible that certain synapses are more susceptible to pruning than others, and this should be addressed in the future. We cannot exclude the possibility that REV-ERBα deletion causes neuronal damage, which elicits microglial phagocytosis of dysfunctional synapses. Indeed, neuronal *C4b* upregulation could be a damage signal. However, the absence of neuronal cell body loss in the dentate gyrus of RKO mice shows that there is no overt neurodegenerative response. These dentate gyrus granule cells give rise to the mossy fiber boutons in the CA3 (Scharfman and Myers, 2012), which are clearly decreased in RKO mice. Thus, the effect of REV-ERBα appears to be specific to the synapses.

Herein, we establish REV-ERBα as a regulator of microglial synaptic phagocytosis. However, we have previously reported that time-of-day changes in microglial morphology were abrogated by REV-ERBα deletion (Griffin et al., 2019). Herein, we observed that microglia engulfed more synapses in the CA3 of the hippocampus at 11PM (ZT17) than at 11AM (ZT5). This was evident by both immunofluorescence and by electron microscopy. To further confirm this striking finding, we used a viral approach in which the retrospenial cortex of mice with GFP labeled microglia had neurons labeled with an mCherry expressing virus. In this paradigm, microglia engulfed more mCherry at 11PM (ZT17) than at 11AM (ZT5). Finally, REV-ERBα deletion abrogated this time-of-day variation in synaptic protein phagocytosis, as RKO mice had high levels of phagocytosis at both timepoints. This data parallels our previous findings with microglial morphologic changes and suggests that REV-ERBα mediates daily changes in the degree of microglial synaptic engulfment. Also, REV-ERBα deletion locks the system in one of the extremes of this naturally occurring oscillation.

Several studies have illustrated the complexity of the interplay between cellular rhythms and sleep in regulating synapses in the brain. Robust circadian rhythms in synaptic protein expression and phosphorylation have bene described which are sleep-dependent (Brüning et al., 2019; Noya et al., 2019). Furthermore, chronic sleep deprivation has also been linked to astrocytic synaptic phagocytosis and microglial activation (Bellesi et al., 2017). This suggests an important role for sleep, which should be explored in future studies. Notably, RKO have subtle alterations in sleep but show grossly normal circadian rhythms in activity and are not sleep deprived (Mang et al., 2016). A study in the rat prefrontal cortex found that synaptic elimination was higher at ZT0 (Choudhury et al., 2020). Since these studies were done in the prefrontal cortex, they suggest potential brain region specific pruning patterns by time of day. Heterogeneity in synaptic pruning has already been described, with areas such as the cerebellum exhibiting greater microglial phagocytic capacity (Ayata et al., 2018). Future work will have to explore the patterns in microglial phagocytosis across more brain regions.

This work has clear implications for neurodegenerative and neuropsychiatric diseases, which have been linked to circadian disruption as well as complement dysregulation and synapse loss. Since REV-ERBα is a nuclear receptor with available small-molecule ligands (Solt et al., 2012), our findings suggest that it could be a therapeutic target for neurological and psychiatric disease. In previous studies in the brain, activation of REV-ERBs appears to suppress microglial cytokine production (Griffin et al., 2019) while inhibition of REV-ERBs can induce microglial amyloid-beta uptake and decrease plaque burden in mice (Lee et al., 2020). Effects on synaptic engulfment should be considered as REV-ERB-based therapeutics are developed.

## MATERIALS AND METHODS

### Mice

Rev-erbα^+/-^ (*Nr1d1*^+/-^) mice on C57bl/6 background were obtained from The Jackson Laboratory (Bar Harbor, ME) and bred at at Washington University. Heterozygous mice were bred together to generate Rev-erbα^+/+^ (wt) and Rev-erbα^-/-^ (RKO) littermates which were used for experiments. Constitutive *Bmal1*^-/-^(BMKO), CAG-Cre^ERT2^, *Aldh1L1*-Cre^ERT2^, *Cx3cr1*-Cre^ERT2^, and *Bmal1*^fl/fl^ mice were obtained from the Jackson Laboratory, and bred so that mice were heterozygous for Cre and homozygous for the floxed *Bmal1* allele. In all experiments, Cre-;Bmal1^f/f^ littermates were used as controls. All inducible knockout lines were were treated with tamoxifen (Sigma) dissolved in corn oil via oral gavage, 2.5mg/day for 5 days, at 2mo. Camk2a-iCre+;Bmal1^fl/fl^ mice were bred at University of Texas Southwestern. CX3CR1^GFP^ mice were obtained from The Jackson Laboratory. Mice were housed on a 12/12 light/dark cycle and fed ad libitum. All procedures performed on the mice were approved by the Washington University IACUC.

### Immunohistochemistry

The following primary antibodies were used (with dilution): Gfap (Rabbit polyclonal, 1:2500, Dako/Agilent Cat# Z0334), C3 (Rat monoclonal, 1:500, Novus Biologicals Cat# NB200-540), Iba1 (Goat polyclonal, 1:500, Abcam Cat# ab5076), Synaptophysin (Mouse monoclonal, 1:100, Abcam Cat# ab8049), CD68 (Rat monoclonal, 1:250, BioRad, Cat# MCA1957), Homer1 (Rabbit polyclonal, 1:500, Synaptic Systems Cat# 160 003), Synaptoporin (Rabbit polyclonal, 1:1000, Synaptic Systems Cat# 102 002), NeuN (Mouse monoclonal, 1:1000, EMD/Millipore Cat# 14-5698-82).

Mice were anesthetized with intraperitoneal (i.p.) injection of pentobarbital (150mg/kg), followed by pump perfusion for 3 mins with ice cold Dulbecco’s modified Phosphate Buffered Saline (DPBS) containing 3g/l heparin sulfate. One hemisphere was drop fixed in 4% paraformaldehyde (PFA) for 24 hours at 4°C, then cryoprotected with 30% sucrose in PBS also at 4°C for at least 48 hours. Brains were embedded in OCT and frozen in acetone with dry ice. 12μm serial coronal sections were cut on a cryostat and mounted directly onto the glass slides for synaptic pruning analysis or synaptic quantification. Sections were washed 3 times in Tris buffered saline (TBS), blocked for 45 mins in TBS containing 20% goat (or donkey) serum, 2% Mouse-on-mouse (M.O.M) blocking reagent and 0.4% Triton X-100 (Sigma-Aldrich, St. Louis, MO). Sections were then incubated in TBS containing 10% goat (or donkey) serum, 8% M.O.M protein concentrate and 0.4% Triton X-100 with primary antibody overnight at 4°C. Sections were then washed 3 times and incubated for 4 hours at room temperature with 1:1000 fluorescent secondary antibody in a solution of TBS containing 10% goat (or donkey) serum, 8% M.O.M protein concentrate and 0.4% Triton X-100. For other non-synaptic staining, 50μm serial coronal sections were cut on a freezing sliding microtome and stored in cryoprotectant (30% ethylene glycol, 15% sucrose, 15% phosphate buffer in ddH_2_O). Sections were washed 3 times in Tris buffered saline (TBS), blocked for 30 mins in TBS containing 3% goat serum and 0.25% Triton X-100 (Sigma-Aldrich, St. Louis, MO) then incubated in TBS containing 1% goat serum and 0.25% Triton X-100 with primary antibody overnight at 4°C. Sections were then washed 3 times and incubated for 1 hour at room temperature with 1:1000 fluorescent secondary antibody and mounted on slides. Confocal images were taken on the Nikon Elements software on the Nikon A1Rsi scanning confocal microscope. Z-stacks were taken at a step size of 0.5-1µm from dark to dark through the tissue.

### Microarray analysis

5-6 mo Rev-erbα^-/-^ and WT littermates were placed in constant darkness for 24 hours and then harvested with i.p. injection of pentobarbital in the dark, following pump perfusion. RNA was isolated from flash frozen hippocampus samples as above and submitted to the Genome Technology Access Center at Washington University for quality control, MessageAmp RNA library preparation and Agilent 4×44k mouse microarray. Raw data was normalized and analyzed using Partek Genomics suite v6.6. For Gene Ontology (GO) term analysis, a list of all genes which were upregulated at least 2-fold in KO with an uncorrected P value <0.05 by 2-tailed T-test were uploaded to DAVID v6.8 (https://david.ncifcrf.gov/). Functional annotation analysis was performed for GO terms related to biological processes. Datasets are freely available on the Array Express Server: https://www.ebi.ac.uk/arrayexpress/experiments/EMTAB-7590

### Quantitative PCR

Flash-frozen brain tissue was homogenized with a mechanical handheld homogenizer for 20 seconds in RNA kit lysis buffer (PureLink™ RNA Mini Kit, Life Technologies, Carlsbad, CA) plus 1% β-mercaptoethanol. RNA was then purified using the kit protocol. Cells well collected and lysed in Trizol (Life Technologies). The aqueous layer was collected following chloroform extraction (added at 1:5 then spun at 13000xg for 15 minutes) with RNA isolation protocol. RNA concentrations were then measured using the Nanodrop spectrophotometer and cDNA was made using a high capacity RNA-cDNA reverse transcription kit (Applied Biosystems/LifeTechnologies) with 1μg RNA used per 20μL reaction. Real-time quantitative PCR was performed with Taqman primers and PCR Master Mix buffer (Life Technologies) on the ABI StepOnePlus 12k Real-Time PCR thermocyclers. β-actin (*Actb*) mRNA levels were used for normalization during analysis. The following primers were used (all from Life Technologies, assay number is listed): *Actb*: Mm02619580_g1, *Nr1d1*: Mm00520708_m1, *C4b*: Mm00437893_g1, *C3*: Mm01232779_m1, *Fabp7*: Mm00437838_m1.

### Electron microscopy

Animals were perfused with a fixative mix consisting of 2.5% glutaraldehyde + 2% paraformaldehyde (fresh, EM grade) in 0.15M cacodylate buffer with 2mM CaCl2 (final concentrations). Brains were extracted, blocked, and 100 µm coronal sections were made using a Leica 1200S vibratome. Tissues were then washed 3 × 10 minutes in cold cacodylate buffer containing 2mM calcium chloride, and then incubated in a solution of 1% OsO_4_ containing 3% potassium ferrocyanide in 0.3M cacodylate buffer with 4mM calcium chloride for 1 hour in the dark. Following incubation, tissues were incubated for 20 minutes in a 1% thiocarbohydrazide (TCH) solution, rinsed 3 × 10 minutes in ddH2O at room temperature and thereafter placed in 2% osmium tetroxide (NOT osmium ferrocyanide) in ddH20 for 30 minutes, at room temperature. Tissues were washed 3 × 10 minutes at room temperature in ddH2O then placed in 1% uranyl acetate (aqueous) and incubated at 4° overnight. Tissues were incubated in a lead aspartate solution at 60°C oven for 30 minutes, washed 5 × 3 minutes in ddH2O, and returned to the lead aspartate solution at 60°C for 30 minutes. Tissues were washed 3 × 10 minutes in room temperature ddH2O and dehydrated in 50%, 70%, 90%, 100%, 100% acetone (anhydrous), 10 minutes each. Samples were embedded in Durcupan ACM resin and polymerized 60°C oven for 48 hours. 70 nm slices were made using a Leica UC7 ultramicrotome, and sections were picked up on a Si wafer (Ted Pella, Redding, CA). Images were acquired on a Zeiss Merlin FE-SEM using a solid state backscatter detector (8kV, 900pA) at 7nm resolution with 5µs pixel dwell times and 4x line averaging. Large area scans of ∼150µm × 150 µm field of view were acquired and stitched using Atlas 5.0 (Fibics, Ottawa, Canada)

### Stereotactic surgery and Intracortical viral Injections

CX3CR1^GFP^ mice had their heads shaved and ear bars put in bilaterally. Iodine was then applied and the skin over the skull was cut open gently using surgical scissors. The skull was checked to make sure that it was level on both sides of the midline. A hole was drilled through the skull but not into the brain parenchyma. The coordinates used for the retrospenial cortex were 0.3mm mediolateral (M/L), −2mm anterior-posterior(A/P) and −1mm dorso-ventral(D/V) with bregma as a reference. The needle was placed in the target location and then allowed to rest for 2 mins before infusion. The AAV2-CaMKII-mCherry virus was obtained from the UNC viral vector core and infused at a rate of 0.2µL/min. In total, the mice were injected with 2μL in the retrospenial cortex. After the infusion, we waited 5 mins for the virus to diffuse in the parenchyma and then the needle was slowly removed. The skin over the skull was then stitched up and antibiotic was applied to the area. The mice were then allowed to recover in an empty cage on a heating pad. After the surgery the mice were checked twice daily, 4 hours apart to ensure survival for 3 days. The mice were then allowed to age for 1 month before being sacrificed for sectioning and imaging.

### Synaptic volume and engulfment analysis/3D reconstructions

Imaris visualization and analysis software (Version 9, Bitplane, South Windsor, CT, USA) was used at the Washington University Center for Cellular Imaging. For all analyses Z-stacks were saved in the .nd2 file format and loaded into the software. 3-D surfaces had a surface detail ranging from 0.1-0.3μm. To quantify the volume of the synapses, we generated 3-D surfaces from each of the synaptic markers (Synaptophysin, Synaptoporin and Homer1). We used the Batch colocalization function to colocalize the Synaptophysin and Homer1 volumes. The total volume for each Z-stack was summed. For the synaptic engulfment analyses, we first generated volumes from synaptophysin, CD68 and Iba1 staining. We then colocalized the synaptophysin volume with the CD68 volume. The resulting volume was then colocalized with the Iba1 volume to produce a final microglial synaptic engulfment volume. All values were normalized to the average values for the control group in each experiment.

### Primary Cell Cultures

WT Neuronal and astrocyte cultures were obtained from CD1 mice from Charles River Laboratories (Wilmington, MA). Neurons were isolated from E16-E18 pups and astrocytes were isolated from P0-2 pups. For experiments involving Rev-erbα^-/-^ astrocytes or microglia, the cultures were made from mice on a C57BL/6J background. Cortices plus hippocampus were dissected and stripped of meninges in ice-cold DMEM (Life Technologies) and then incubated in 0.05% Trypsin-EDTA at 37°C for 15 mins (Neurons) or 20 mins (Astrocytes). Tissue was gently triturated in 37°C DMEM plus 10% FBS (Gibco). For astrocyte cultures cells were then plated in T75 flasks coated for 2 hours at 37°C with 50µg/mL poly-D-Lysine (PDL – MP Biosciences, Santa Ana, CA) then rinsed with ddH_2_O. Astrocytes were then grown to confluence in DMEM, 10% FBS, and 1% Penicillin/Streptomycin (P/S) with or without 5 ng/mL GM-CSF. For primary microglia isolation from GM-CSF containing media, flasks were shaken at 225rpm at 37°C for 2 hours and replated on plates coated with PDL. For neuronal cultures, triturated cells were transferred to a second tube to remove debris, then diluted in Neurobasal (Life Technologies) plus B27 (Life Technologies) prior to plating on a bed of astrocytes or a PDL-coated plate. For co-culture experiments, astrocytes were grown to confluence following replating in 24-well plates and neurons were added immediately after primary dissection at a concentration of 150,000 cells/mL in Neurobasal plus B27 supplement and glutamine (Life Technologies). After 48 hours, 50% of the media was changed to Neurobasal plus anti-oxidant free B-27 (AOF B-27, Life Technologies) and glutamine. Following that, 50% media changes were done using Neurobasal plus AOF B-27 and glutamine every 3 days until neuronal DIV 11 at which the experimental wells received H_2_O_2_ and the controls received vehicle for 24 hours. All wells were then fixed with 4% PFA and stained. In general, all cell culture experiments were performed in duplicate and repeated at least 3 times, using primary cells prepared from at least separate dissections

### Statistical analysis

Statistical analyses were performed using GraphPad Prism 8. We performed t-tests with Welch’s correction for multiple comparisons.

## ACKNOWLEDGEMENTS

This work was supported by awards from the Cure Alzheimer’s Fund (ESM), Coins for Alzheimer’s Research Trust (ESM), and NIH grants R01AG054517 and R01AG063743 (ESM). PG was supported by National Science Foundation Grant DGE-1745038. Imaging was performed with support from the Washington University Center for Cellular Imaging (WUCCI), which is funded by Washington University, Children’s Discovery Institute (CDI-CORE-2015-505), and the Foundation for Barnes-Jewish Hospital (3770). The authors thank Drs. Mariko Izumo and Joseph Takahashi (UT Southwestern Medical School) for providing brain tissue samples from Camk2a-iCre;Bmal1(f/f) mice.

## REFERENCES

1. Ayata, P., Badimon, A., Strasburger, H.J., Duff, M.K., Montgomery, S.E., Loh, Y.E., Ebert, A., Pimenova, A.A., Ramirez, B.R., Chan, A.T., et al. (2018). Epigenetic regulation of brain region-specific microglia clearance activity. Nature neuroscience 21, 1049–1060.

2. Bellesi, M., de Vivo, L., Chini, M., Gilli, F., Tononi, G., and Cirelli, C. (2017). Sleep Loss Promotes Astrocytic Phagocytosis and Microglial Activation in Mouse Cerebral Cortex. J Neurosci 37, 5263–5273.

3. Braak, H., Alafuzoff, I., Arzberger, T., Kretzschmar, H., and Del Tredici, K. (2006). Staging of Alzheimer disease-associated neurofibrillary pathology using paraffin sections and immunocytochemistry. Acta neuropathologica 112, 389–404.

4. Brüning, F., Noya, S.B., Bange, T., Koutsouli, S., Rudolph, J.D., Tyagarajan, S.K., Cox, J., Mann, M., Brown, S.A., and Robles, M.S. (2019). Sleep-wake cycles drive daily dynamics of synaptic phosphorylation. Science 366, eaav3617.

5. Choudhury, M.E., Miyanishi, K., Takeda, H., Islam, A., Matsuoka, N., Kubo, M., Matsumoto, S., Kunieda, T., Nomoto, M., Yano, H., et al. (2020). Phagocytic elimination of synapses by microglia during sleep. Glia 68, 44–59.

6. Chung, S., Lee Eun J., Yun, S., Choe Han K., Park, S.-B., Son Hyo J., Kim, K.-S., Dluzen Dean E., Lee, I., Hwang, O., et al. (2014). Impact of Circadian Nuclear Receptor REV-ERBα on Midbrain Dopamine Production and Mood Regulation. Cell 157, 858–868.

7. Everett, L.J., and Lazar, M.A. (2014). Nuclear receptor Rev-erbalpha: up, down, and all around. Trends in endocrinology and metabolism: TEM 25, 586–592.

8. Fonken, L.K., Frank, M.G., Kitt, M.M., Barrientos, R.M., Watkins, L.R., and Maier, S.F. (2015). Microglia inflammatory responses are controlled by an intrinsic circadian clock. Brain Behav Immun 45, 171–179.

9. Griffin, P., Dimitry, J.M., Sheehan, P.W., Lananna, B.V., Guo, C., Robinette, M.L., Hayes, M.E., Cedeno, M.R., Nadarajah, C.J., Ezerskiy, L.A., et al. (2019). Circadian clock protein Rev-erbalpha regulates neuroinflammation. Proc Natl Acad Sci U S A 116, 5102–5107.

10. Grosse, G., Tapp, R., Wartenberg, M., Sauer, H., Fox, P.A., Grosse, J., Gratzl, M., and Bergmann, M. (1998). Prenatal hippocampal granule cells in primary cell culture form mossy fiber boutons at pyramidal cell dendrites. Journal of Neuroscience Research 51, 602–611.

11. Harding, H.P., and Lazar, M.A. (1995). The monomer-binding orphan receptor Rev-Erb represses transcription as a dimer on a novel direct repeat. Mol Cell Biol 15, 4791–4802.

12. Hayashi, Y., Koyanagi, S., Kusunose, N., Okada, R., Wu, Z., Tozaki-Saitoh, H., Ukai, K., Kohsaka, S., Inoue, K., Ohdo, S., et al. (2013). The intrinsic microglial molecular clock controls synaptic strength via the circadian expression of cathepsin S. Sci Rep 3, 2744.

13. Hong, S., Beja-Glasser, V.F., Nfonoyim, B.M., Frouin, A., Li, S., Ramakrishnan, S., Merry, K.M., Shi, Q., Rosenthal, A., Barres, B.A., et al. (2016). Complement and microglia mediate early synapse loss in Alzheimer mouse models. Science (New York, NY 352, 712–716.

14. Izumo, M., Pejchal, M., Schook, A.C., Lange, R.P., Walisser, J.A., Sato, T.R., Wang, X., Bradfield, C.A., and Takahashi, J.S. (2014). Differential effects of light and feeding on circadian organization of peripheral clocks in a forebrain Bmal1 mutant. Elife 3.

15. Jung, S., Aliberti, J., Graemmel, P., Sunshine, M.J., Kreutzberg, G.W., Sher, A., and Littman, D.R. (2000). Analysis of fractalkine receptor CX(3) CR1 function by targeted deletion and green fluorescent protein reporter gene insertion. Mol Cell Biol 20, 4106–4114.

16. Kim, Y.H., Marhon, S.A., Zhang, Y., Steger, D.J., Won, K.J., and Lazar, M.A. (2018). Rev-erbalpha dynamically modulates chromatin looping to control circadian gene transcription. Science 359, 1274–1277.

17. Lananna, B.V., Nadarajah, C.J., Izumo, M., Cedeno, M.R., Xiong, D.D., Dimitry, J., Tso, C.F., McKee, C.A., Griffin, P., Sheehan, P.W., et al. (2018). Cell-Autonomous Regulation of Astrocyte Activation by the Circadian Clock Protein BMAL1. Cell Rep 25, 1–9.e5.

18. Lee, J., Kim, D.E., Griffin, P., Sheehan, P.W., Kim, D.-H., Musiek, E.S., and Yoon, S.-Y. (2020). Inhibition of REV-ERBs stimulates microglial amyloid-beta clearance and reduces amyloid plaque deposition in the 5XFAD mouse model of Alzheimer’s disease. Aging Cell 19, e13078–e13078.

19. Lehrman, E.K., Wilton, D.K., Litvina, E.Y., Welsh, C.A., Chang, S.T., Frouin, A., Walker, A.J., Heller, M.D., Umemori, H., Chen, C., et al. (2018). CD47 Protects Synapses from Excess Microglia-Mediated Pruning during Development. Neuron 100, 120–134.e126.

20. Lian, H., Yang, L., Cole, A., Sun, L., Chiang, A.C., Fowler, S.W., Shim, D.J., Rodriguez-Rivera, J., Taglialatela, G., Jankowsky, J.L., et al. (2015). NFkappaB-activated astroglial release of complement C3 compromises neuronal morphology and function associated with Alzheimer’s disease. Neuron 85, 101–115.

21. Litvinchuk, A., Wan, Y.W., Swartzlander, D.B., Chen, F., Cole, A., Propson, N.E., Wang, Q., Zhang, B., Liu, Z., and Zheng, H. (2018). Complement C3aR Inactivation Attenuates Tau Pathology and Reverses an Immune Network Deregulated in Tauopathy Models and Alzheimer’s Disease. Neuron.

22. Mang, G.M., La Spada, F., Emmenegger, Y., Chappuis, S., Ripperger, J.A., Albrecht, U., and Franken, P. (2016). Altered Sleep Homeostasis in Rev-erbalpha Knockout Mice. Sleep 39, 589–601.

23. Mohawk, J.A., Green, C.B., and Takahashi, J.S. (2012). Central and peripheral circadian clocks in mammals. Annual review of neuroscience 35, 445–462.

24. Musiek, E.S., and Holtzman, D.M. (2016). Mechanisms linking circadian clocks, sleep, and neurodegeneration. Science 354, 1004–1008.

25. Musiek, E.S., Lim, M.M., Yang, G., Bauer, A.Q., Qi, L., Lee, Y., Roh, J.H., Ortiz-Gonzalez, X., Dearborn, J.T., Culver, J.P., et al. (2013). Circadian clock proteins regulate neuronal redox homeostasis and neurodegeneration. J Clin Invest 123, 5389–5400.

26. Noya, S.B., Colameo, D., Brüning, F., Spinnler, A., Mircsof, D., Opitz, L., Mann, M., Tyagarajan, S.K., Robles, M.S., and Brown, S.A. (2019). The forebrain synaptic transcriptome is organized by clocks but its proteome is driven by sleep. Science 366, eaav2642.

27. Parkhurst, C.N., Yang, G., Ninan, I., Savas, J.N., Yates, J.R., 3rd, Lafaille, J.J., Hempstead, B.L., Littman, D.R., and Gan, W.B. (2013). Microglia promote learning-dependent synapse formation through brain-derived neurotrophic factor. Cell 155, 1596–1609.

28. Preitner, N., Damiola, F., Lopez-Molina, L., Zakany, J., Duboule, D., Albrecht, U., and Schibler, U. (2002). The orphan nuclear receptor REV-ERBalpha controls circadian transcription within the positive limb of the mammalian circadian oscillator. Cell 110, 251–260.

29. Retnakaran, R., Flock, G., and Giguere, V. (1994). Identification of RVR, a novel orphan nuclear receptor that acts as a negative transcriptional regulator. Molecular endocrinology (Baltimore, Md) 8, 1234–1244.

30. Schafer, D.P., Lehrman, E.K., Kautzman, A.G., Koyama, R., Mardinly, A.R., Yamasaki, R., Ransohoff, R.M., Greenberg, M.E., Barres, B.A., and Stevens, B. (2012). Microglia sculpt postnatal neural circuits in an activity and complement-dependent manner. Neuron 74, 691–705.

31. Scharfman, H.E., and Myers, C.E. (2012). Hilar mossy cells of the dentate gyrus: a historical perspective. Front Neural Circuits 6, 106.

32. Schnell, A., Chappuis, S., Schmutz, I., Brai, E., Ripperger, J.A., Schaad, O., Welzl, H., Descombes, P., Alberi, L., and Albrecht, U. (2014). The nuclear receptor REV-ERBalpha regulates Fabp7 and modulates adult hippocampal neurogenesis. PLoS One 9, e99883.

33. Sekar, A., Bialas, A.R., de Rivera, H., Davis, A., Hammond, T.R., Kamitaki, N., Tooley, K., Presumey, J., Baum, M., Van Doren, V., et al. (2016). Schizophrenia risk from complex variation of complement component 4. Nature 530, 177–183.

34. Selkoe, D.J. (2002). Alzheimer’s disease is a synaptic failure. Science 298, 789–791.

35. Singec, I., Knoth, R., Ditter, M., Hagemeyer, C.E., Rosenbrock, H., Frotscher, M., and Volk, B. (2002). Synaptic vesicle protein synaptoporin is differently expressed by subpopulations of mouse hippocampal neurons. J Comp Neurol 452, 139–153.

36. Solt, L.A., Wang, Y., Banerjee, S., Hughes, T., Kojetin, D.J., Lundasen, T., Shin, Y., Liu, J., Cameron, M.D., Noel, R., et al. (2012). Regulation of circadian behaviour and metabolism by synthetic REV-ERB agonists. Nature 485, 62–68.

37. Stevanovic, K., Yunus, A., Joly-Amado, A., Gordon, M., Morgan, D., Gulick, D., and Gamsby, J. (2017). Disruption of normal circadian clock function in a mouse model of tauopathy. Experimental neurology 294, 58–67.

38. Stevens, B., Allen, N.J., Vazquez, L.E., Howell, G.R., Christopherson, K.S., Nouri, N., Micheva, K.D., Mehalow, A.K., Huberman, A.D., Stafford, B., et al. (2007). The classical complement cascade mediates CNS synapse elimination. Cell 131, 1164–1178.

39. Sulli, G., Rommel, A., Wang, X., Kolar, M.J., Puca, F., Saghatelian, A., Plikus, M.V., Verma, I.M., and Panda, S. (2018). Pharmacological activation of REV-ERBs is lethal in cancer and oncogene-induced senescence. Nature 553, 351–355.

40. Takahashi, J.S. (2017). Transcriptional architecture of the mammalian circadian clock. Nature reviews Genetics 18, 164–179.

41. Videnovic, A., Lazar, A.S., Barker, R.A., and Overeem, S. (2014). ‘The clocks that time us’-- circadian rhythms in neurodegenerative disorders. Nature reviews Neurology 10, 683–693.

42. Welch, R.D., Guo, C., Sengupta, M., Carpenter, K.J., Stephens, N.A., Arnett, S.A., Meyers, M.J., Sparks, L.M., Smith, S.R., Zhang, J., et al. (2017). Rev-Erb co-regulates muscle regeneration via tethered interaction with the NF-Y cistrome. Molecular metabolism 6, 703–714.

43. Werneburg, S., Jung, J., Kunjamma, R.B., Ha, S.K., Luciano, N.J., Willis, C.M., Gao, G., Biscola, N.P., Havton, L.A., Crocker, S.J., et al. (2020). Targeted Complement Inhibition at Synapses Prevents Microglial Synaptic Engulfment and Synapse Loss in Demyelinating Disease. Immunity 52, 167–182.e167.

44. Zhang, Y., Chen, K., Sloan, S.A., Bennett, M.L., Scholze, A.R., O’Keeffe, S., Phatnani, H.P., Guarnieri, P., Caneda, C., Ruderisch, N., et al. (2014). An RNA-sequencing transcriptome and splicing database of glia, neurons, and vascular cells of the cerebral cortex. J Neurosci 34, 11929–11947.

45. Zhang, Y., Fang, B., Emmett, M.J., Damle, M., Sun, Z., Feng, D., Armour, S.M., Remsberg, J.R., Jager, J., Soccio, R.E., et al. (2015). GENE REGULATION. Discrete functions of nuclear receptor Rev-erbalpha couple metabolism to the clock. Science 348, 1488–1492.

46. Zhou, Y., Song, W.M., Andhey, P.S., Swain, A., Levy, T., Miller, K.R., Poliani, P.L., Cominelli, M., Grover, S., Gilfillan, S., et al. (2020). Human and mouse single-nucleus transcriptomics reveal TREM2-dependent and TREM2-independent cellular responses in Alzheimer’s disease. Nat Med 26, 131–142.

47. Zipfel, P.F., and Skerka, C. (2009). Complement regulators and inhibitory proteins. Nature reviews Immunology 9, 729–740.

