## Supplemental Figures for "REV-ERBα mediates complement expression and circadian regulation of microglial synaptic phagocytosis"

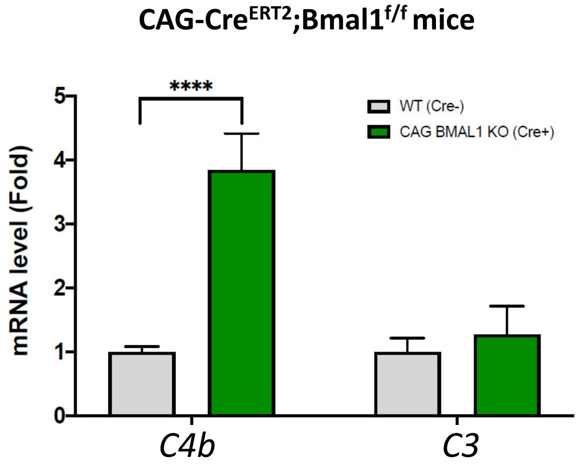


**Figure S1: Post-natal *Bmal1* deletion causes early induction of *C4b*.** qPCR analysis for complement genes in hippocampal tissue from global inducible Bmal1 KO mice (CAG-Cre$^{ERT2}$;Bmal1$^{f/f}$) and Cre- littermate controls. N = 5-6/group. Mice were all treated with tamoxifen at 2mo and harvested at 4mo. ****p<0.00001 by 2-tailed T-test with Welch's correction.

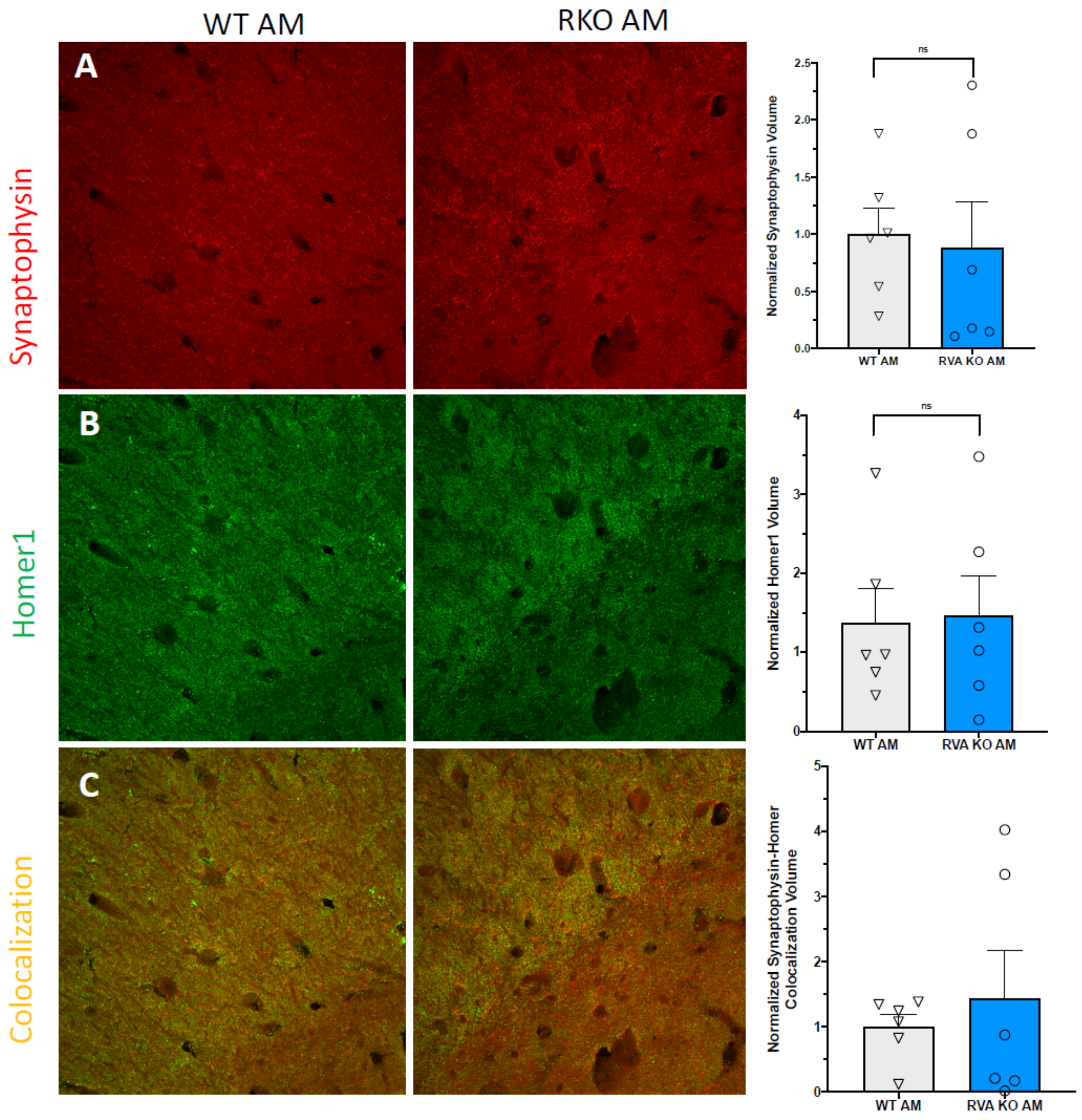


**Figure S2: Rev-erbα deletion does not induce synapse loss in CA1.** A. Representative 60X maximum intensity projects of synaptophysin staining in the CA1 region of WT or RKO mice as well as the associated normalized volume quantifications (N = 5/group). B. Representative 60X maximum intensity projects of Homer1 staining in the CA1 region of WT or RKO mice as well as the associated normalized volume quantifications (N = 5/group). C. Representative 60X maximum intensity projects of Colocalized Synaptophysin and Homer1 staining in the CA1 region of WT or RKO mice as well as the associated normalized volume quantifications (N = 5/group).
